# Reuniens thalamus recruits recurrent excitation in medial prefrontal cortex

**DOI:** 10.1101/2024.05.31.596906

**Authors:** Gil Vantomme, Gabrielle Devienne, Jacob M Hull, John R Huguenard

**Author notes:** **Corresponding author:** John R Huguenard, Stanford Neuroscience Building S189, 290 Jane Stanford Way, Stanford, CA 94305-5088, **Email:**.

## Abstract

Medial prefrontal cortex (mPFC) and hippocampus are critical for memory retrieval, decision making and emotional regulation. While ventral CA1 (vCA1) shows direct and reciprocal connections with mPFC, dorsal CA1 (dCA1) forms indirect pathways to mPFC, notably via the thalamic Reuniens nucleus (Re). Neuroanatomical tracing has documented structural connectivity of this indirect pathway through Re however, its functional operation is largely unexplored. Here we used *in vivo* and *in vitro* electrophysiology along with optogenetics to address this question. Whole-cell patch-clamp recordings in acute mouse brain slices revealed both monosynaptic excitatory responses and disynaptic feedforward inhibition at Re-mPFC synapses. However, we also identified a novel prolonged excitation of mPFC by Re. These early monosynaptic and late recurrent components are in marked contrast to the primarily feedforward inhibition characteristic of thalamic inputs to neocortex. Local field potential recordings in mPFC brain slices revealed prolonged synaptic activity throughout all cortical lamina upon Re activation, with the late excitation enhanced by blockade of parvalbumin neurons and GABA_A_Rs. *In vivo* Neuropixels recordings in head-fixed awake mice revealed a similar prolonged excitation of mPFC units by Re activation. In summary, Re output produces recurrent feedforward excitation within mPFC suggesting a potent amplification system in the Re-mPFC network. This may facilitate amplification of dCA1->mPFC signals for which Re acts as the primary conduit, as there is little direct connectivity. In addition, the capacity of mPFC neurons to fire bursts of action potentials in response to Re input suggests that these synapses have a high gain.

**Significance statement:** The interactions between medial prefrontal cortex and hippocampus are crucial for memory formation and retrieval. Yet, it is still poorly understood how the functional connectivity of direct and indirect pathways underlies these functions. This research explores the synaptic connectivity of the indirect pathway through the Reuniens nucleus of the thalamus using electrophysiological recordings and optogenetic manipulations. The study found that Reuniens stimulation recruits recurrent and long-lasting activity in mPFC - a phenomenon not previously recorded. This recurrent activity might create a temporal window ideal for coincidence detection and be an underlying mechanism for memory formation and retrieval.

## Introduction

The direct and indirect pathways between the hippocampus and the medial prefrontal cortex (mPFC) play a significant role in emotional, contextual and spatial processing (1, 2). The direct pathway is generally associated with working memory, learning, and contextual processing (2). The indirect pathways, through intermediary regions like the thalamic Reuniens nucleus (Re), might be important for controlling the quantity and type of information retrieved (3). Accumulating evidence suggests that the Re plays a central role in coordinating neuronal activity within the mPFC and hippocampus. Lesions (4) or inhibition (5, 6) of Re reduce coherence between mPFC and hippocampus, in particular in the delta (2-5Hz) and gamma (30-90Hz) frequency ranges. Recently, optogenetic activation of Re was shown to increase mPFC/hippocampus beta (15-30Hz) coherence, which is linked with non-spatial sequence memory (7).

Behaviorally, the Re has been linked to many cognitive processes, most notably: goal directed behavior, working memory, inhibitory control, avoidance and behavioral flexibility (for review (8)). Large efforts have been made in recent years to evaluate the behavioral consequences of permanent lesions, optogenetic manipulation, and pharmacological inhibition of Re in rodents. However, the impact of Re lesions in recognition tasks is controversial. Jung et al. (9) reported a reduction in the exploration time of novel object/displaced object while Barker and Warburton did not observe these changes (10). Lesion of Re led to an increase in performance in some attention dependent tasks (11) but deficits in others (12). Inactivation of Re increased generalization of aversive memory (13) and deficit in the performance in tasks requiring spatial working memory (14) or its encoding (15).

The broad spectrum of functions in which Re is implicated and the disparities observed across these studies emphasizes the complexity of the indirect pathways between hippocampus and mPFC. Understanding this network requires an in-depth exploration of the circuit connectivity that underpins its functionality. However, little is known about the synaptic properties, neurotransmitter receptor content, plasticity profile or synaptic integration at the Re-mPFC.

To address this issue, the present study combined optogenetic activation of Re afferents with electrophysiological recordings of its postsynaptic targets both in vitro and in vivo. In vitro whole-cell patch-clamp recordings combined with pharmacological approaches provided detailed functional properties of Re-mPFC synapses and unmasked a strong feedforward excitatory dynamic specific to mPFC. In vitro local field potential recordings in mPFC further confirmed the presence of this recurrent excitation of the mPFC network. Neuropixels probe single unit recordings in awake behaving animals revealed that activation of Re leads to prolonged firing in mPFC, recapitulating observations made in vitro.

Altogether, these data show the functional synaptic connectivity, short-term plasticity profile, and neurotransmitter receptor expression that underlie the function of the indirect pathway between hippocampus and mPFC through Re. They also show how the mPFC network uniquely amplifies Re output, which may be one of the mechanisms underlying modulation of memory retrieval.

## Results

### Re activation induces prolonged excitation in mPFC neurons

Whole-cell patch-clamp recordings of mPFC neurons were obtained from acute brain slices from mice injected in Re with AAV1-CaMKIIa-ChR2-eYFP (Fig 1A). eYFP/ChR2 positive Re axons in mPFC showed expression in layer 1 and deep layers, consistent with previous studies (3, 16). Activation of Re afferents induced excitatory postsynaptic currents (EPSCs) in mPFC neurons across all layers (Fig 1B). Beyond classical monophasic EPSCs, evident by a single peak, a fast rise and exponential decay (Fig 1B, black), about half of L2/3 and L5 pyramidal cells and 1/5 L1 interneuron in mPFC showed compound EPSCs with a prominent late peak (Fig 1B, blue and Fig 1E). To test whether such EPSC amplification was specific to the Re-mPFC pathway, we compared the response to that obtained in L5 pyramidal cells in entorhinal cortex (EC), another cortical target of Re (Fig 1C). In contrast to mPFC, EC neurons only responded with classical monophasic EPSCs upon Re activation (Fig 1D, Suppl. Fig 1). Late EPSCs in mPFC were defined as having a prominent peak in their first derivative with a rate of change and minimum peak prominence of at least 0.1 pA/ms, 10 ms or more after light onset (Fig 1F, Wilcoxon rank sum test, latency of monophasic vs late EPSC: p=9×10^−10^). This classification revealed that a significant proportion of L2/3 and L5 pyramidal cells showed these late EPSCs upon Re activation compared to EC (Fig 1E, Chi-square tests, L2/3-mPFC vs L5-EC: p=0.005, L5-mPFC vs L5-EC: p=0.0001). A complementary approach to assess contributions of late EPSCs to the overall Re-induced response is to measure the weighted decay time constant (τ_w_), which provides an objective measure of response duration (Suppl Fig 2A). The EPSC τ_w_ is longer in L2/3 and L5 pyramidal cells than in EC neurons (Suppl. Fig 2B). The proportion of cells with an EPSC τ_w_ above 30 ms is similar to the proportion of cells with a prominent late EPSC (Suppl. Fig 2C). These late EPSCs likely reflect prolonged and/or polysynaptic activity in mPFC. To test for this, we determined whether late events were polysynaptic. Accordingly, in a subset of 7 mPFC L5 pyramidal cells, we blocked the light-evoked EPSCs with the sodium channel blocker, tetrodotoxin (TTX) (Wilcoxon-signed rank test, ACSF vs TTX: p=0.016), and partially recovered them in all cells by bath application of the potassium channel blocker, 4-aminopyridine (4AP) (Wilcoxon-signed rank tests, TTX vs 4AP: p=0.016, ACSF vs 4AP: p=0.016), confirming that at least part of the response is mediated through a monosynaptic connection (17) (Fig 1GH). Late EPSCs were present in 3/7 of these cells and while short-latency responses did recover with 4AP bath application, the late responses did not, indicating that they are mediated through a polysynaptic mechanism within the mPFC. These data strongly suggest that mPFC uniquely amplifies Re inputs through recurrent excitation of the local network. This phenomenon was absent at Re-EC connection and rare at the synaptic connection between posterior thalamus (PO) and L5 pyramidal cells in primary somatosensory cortex (S1), another higher order thalamocortical connection with thalamic axons targeting the apical dendrite in layer 1 (Suppl. Fig 3).

**Figure 1.**
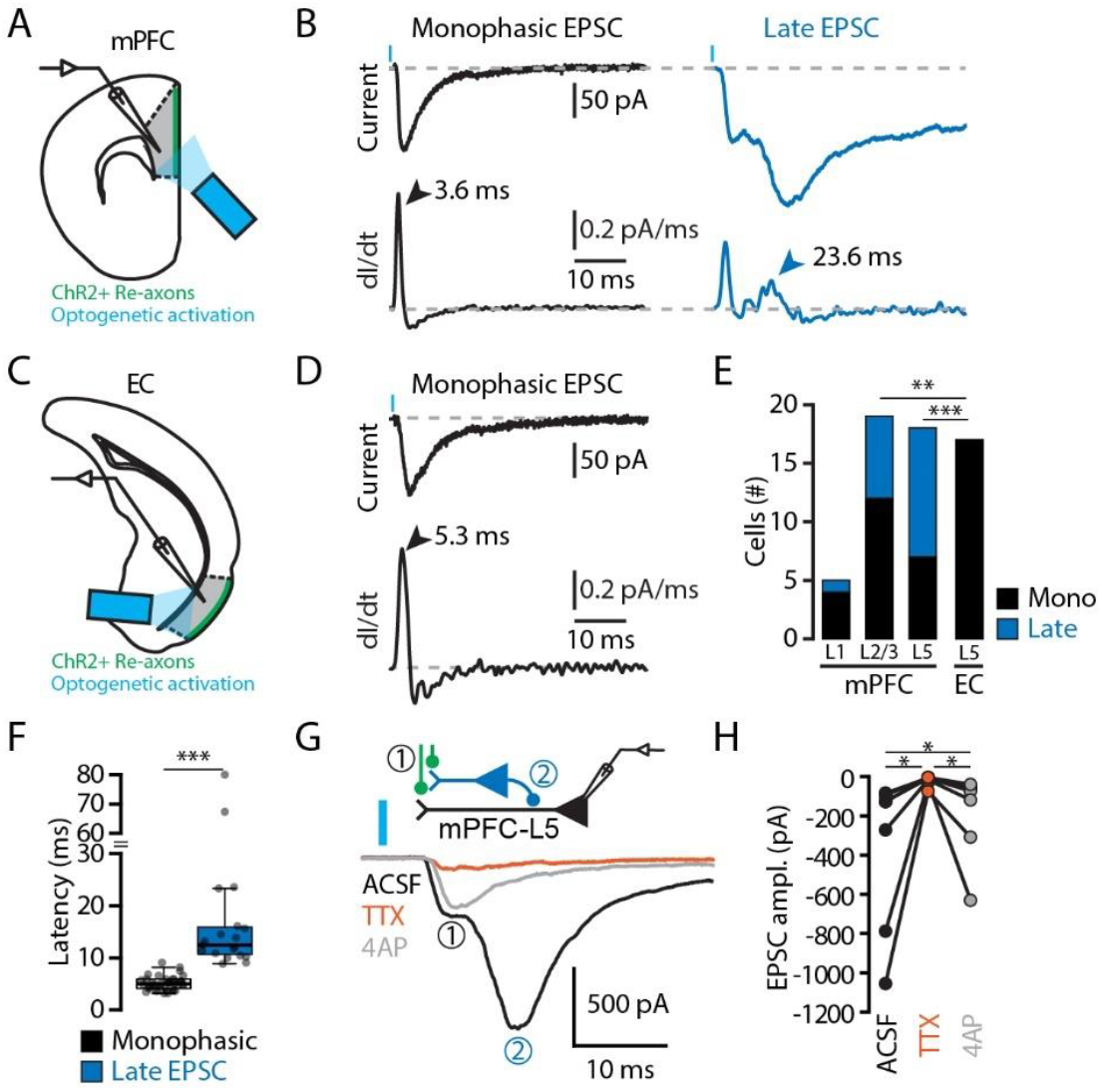
Re activation induces prolonged excitation in mPFC neurons. (A) Experimental approach: whole-cell patch-clamp recording of mPFC neurons combined with optogenetic stimulation of Re axons. (B) Typical traces showing EPSCs (top) after light activation of Re afferents and their corresponding first derivative traces (bottom). Classical monophasic EPSCs showed a single prominent peak in the first derivative (left, black) while complex EPSCs also showed a late (>10 ms after light onset) prominent peak (right, blue). (C) Same as A in EC. (D) Same as B in EC. (E) Histogram of the number of cells with monophasic EPSC (black) or late EPSC (blue) across mPFC layers and in L5 EC. (F) Box-and-whisker plot of the latency of the monophasic EPSCs and late EPSCs from light onset. (G) Top, schematic illustrating Re recruitment of feedforward excitation. Bottom, example traces of an EPSC in ACSF (black), after TTX (orange) and 4AP (grey) bath application. (H) EPSC amplitudes in ACSF (black), TTX (orange) and 4AP (grey).

### Re-dependent late synaptic currents in mPFC are mediated by excitatory synapses

Thalamocortical inputs classically recruit polysynaptic feedforward inhibition (18, 19). The late synaptic currents recorded in mPFC shown in figure 1 are unlikely to be mediated through GABA receptors because with our recording conditions of holding potential at -70 mV, near the reversal potential for Cl^−^, there is little driving force to support GABAergic signaling. Changing the holding potential revealed that Re inputs to mPFC recruit both monosynaptic excitation and polysynaptic responses, with the latter showing both inhibitory and excitatory components (Fig 2AB). Under these conditions, we hypothesized that disinhibition, i.e. blockade of GABAergic signaling, would result in a further increase in the late EPSCs. At -70 mV, the early EPSCs were time-locked to the light activation of Re axons and showed short latencies (4.4+/-0.3 ms) and sub millisecond jitter (0.36+/-0.05 ms). At +10 mV IPSCs showed a longer latency (IPSC: 7.8+/-0.4 ms; EPSC: 4.0+/-0.3 ms), consistent with a classical multisynaptic, feedforward inhibitory circuit (Wilcoxon signed-rank test, EPSC vs IPSC latency: p=4.9×10^−4^). Bath application of gabazine, a GABA_A_ receptor blocker, fully blocked the IPSC (Fig 2BC, ACSF: 229+/-41 pA; Gabazine: 10+/-4 pA; Paired Student *t* test, p=0.002) and also resulted in either the new appearance or increase of the late EPSC (Fig 2B,D, Synaptic charge, ACSF: -4+/-1 pC; Gabazine: -85+/-65 pC; Wilcoxon signed-rank test, p=0.03). Interestingly, activation of Re consistently recruited feedforward inhibition across all mPFC layers, including in L1 interneurons (Suppl. Fig 4AB), leading to an average Excitation/Inhibition ratio below 1 (mPFC-L5: 0.6+/-0.1, n=10; mPFC-L2/3: 0.8+/-0.2, n=6; mPFC-L1: 0.8+/-0.2, n=5). By contrast, Re recruitment of feedforward inhibition in EC L5 neurons was much weaker (EC-L5 E/I ratio: 5.9+/-1.8, n=20; Suppl. Fig 4AB), consistent with the idea that Re inputs more powerfully recruit polysynaptic recurrent activity in mPFC than in EC.

**Figure 2.**
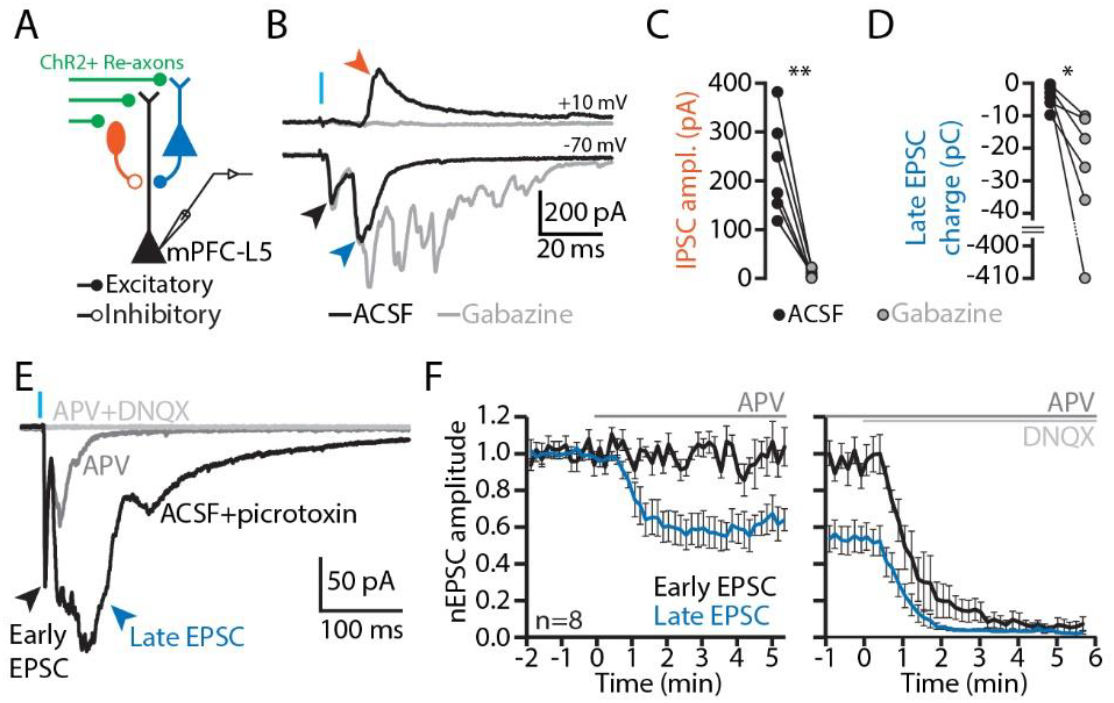
Re-dependent late synaptic currents in mPFC are mediated by excitatory synapses. (A) Schematic illustrating the experimental approach and the recruitment of neurons for feedforward inhibition (orange) and excitation (blue) and the recorded cell (black). (B) Example light evoked EPSC recorded at - 70 mV (bottom) and IPSC recorded at +10 mV (top) in ACSF (black) and in gabazine (grey). Note the presence of two EPSCs, one early within 5 ms of light stimulus, and a late compound EPSC, after a delay of about 10-15 ms. (C) IPSC amplitude in ACSF (black) and gabazine (grey). (D) Charge of the late EPSC in ACSF (black) and gabazine (grey). (E) Example traces of EPSCs recorded at -70 mV in ACSF (black) and after APV (dark grey) and DNQX (light grey) bath application. (F) Effects of APV (left) and DNQX (right) on the amplitude of the early (black) and late (blue) components of the EPSC. The EPSC amplitude was normalized to the amplitude during the baseline. Significance: * p < 0.05, ** p < 0.01, and *** p < 0.001.

Blockade of GABA_A_Rs with picrotoxin permitted stable recording of these late EPSCs. Under these conditions, addition of APV, an NMDAR blocker, strongly reduced the amplitude, but did not eliminate, the late EPSCs (Fig 2EF). Subsequent bath application of DNQX completely abolished the late component (Friedman’s test, p=3.4×10^−4^, post hoc Wilcoxon signed-rank tests, BL vs APV: p= 0.008, APV vs DNQX: p=0.008, BL vs DNQX: p=0.008). By contrast, the early, presumed monosynaptic, component of the EPSC showed little to no reduction in its amplitude after bath application of APV and was fully blocked after subsequent application of DNQX (Friedman’s test, p=8.1×10^−4^, post hoc Wilcoxon signed-rank tests, BL vs APV: p= 0.11, APV vs DNQX: p=0.008, BL vs DNQX: p=0.008). The light-evoked EPSCs showed a NMDAR/AMPAR ratio of 0.21+/-0.03 (Suppl. Fig4A), with a significant contribution from GluN2B containing NMDARs (Suppl. Fig 4B), and a short-term depression profile (Table 1) that is consistent with a high probability of release as shown in previous reports in rat (20).

Altogether, these data show that activation of Re recruits a direct monosynaptic excitatory connection to mPFC neurons as well as both a classical feedforward inhibition and a novel prominent feedforward excitation that to our knowledge has not previously been reported. The timing of the polysynaptic feedforward inhibition (onset 11.9+/-1.9 ms from the light stimulus, n=12) and polysynaptic feedforward excitation (onset 18.9+/-3.2 ms, n=30) are similar, suggesting that the shunting effect of inhibitory synapses would reduce/occlude the late EPSCs. This is consistent with the finding that pharmacological blockade of GABA_A_Rs unmasked the feedforward excitation in the mPFC network.

### Local field potential recordings in mPFC slices revealed prolonged synaptic activity upon Re optogenetic activation

We next examined the overall mPFC network response to Re activation, to test whether we would see further evidence of late network activation that would support the polysynaptic activity observed with whole-cell recordings. We recorded local field potentials (LFPs) in mPFC using a 16-shank linear electrode array spanning across cortical layers (Fig 3A). Slices were collected from mice expressing ChR2 in Re axons following injection with AAV1-CaMKIIa-ChR2-eYFP in Re. Two sets of mice were used, either C57BL6/J or parvalbumin (PV)-cre mice. With the latter an AAV8-hSyn-DIO-hM4D(Gi)-mCherry virus was injected in mPFC to express the inhibitory DREADD in PV neurons (Fig 3BC). In either case, blue light activation of ChR2 in Re afferents resulted in LFPs across mPFC layers. From these we derived the current source density (CSD) (21, 22) (Fig 3DE). In the CSD signals, pairs of sinks and sources were observed following optogenetic stimulation of Re afferents, attributed to cations flowing from the extracellular space into the cells during synaptic excitation, and flowing back out of cells at a distance, respectively. Similarly, during synaptic inhibition, current sources are seen at locations of inhibitory synapses where chloride ions flow into cells, and sinks at other sites where current flows back out of cells. Sources and sinks were prominent within the first 10 ms after the light onset in L2/3 and L5 (Fig 3D). The peak CSD sinks (L2: -31+/-6 µA/mm^3^, L5: -18+/-2 µA/mm^3^, n=9) and sources (L2: 33+/-6 µA/mm^3^, L5: 18+/-4 µA/mm^3^, n=9) were higher in L2/3 than in L5 (Sinks: Paired Student’s *t* test, p=0.03, Sources: Wilcoxon signed rank test, p=0.01), suggesting a stronger synaptic recruitment of L2/3 by Re afferents. To assess polysynaptic activity, we next calculated the peak CSD signals in a window (10 to 30 ms after light onset) that would exclude monosynaptic responses (c.f. Fig. 1G). Both L2/3 and L5 showed sinks (L2: -10+/-2 µA/mm^3^, L5: -11+/-2 µA/mm^3^) and sources (L2: 18+/-3 µA/mm^3^, L5: 10+/-3 µA/mm^3^), which likely reflect a combination of disynaptic feedforward inhibition and excitation.

**Figure 3.**
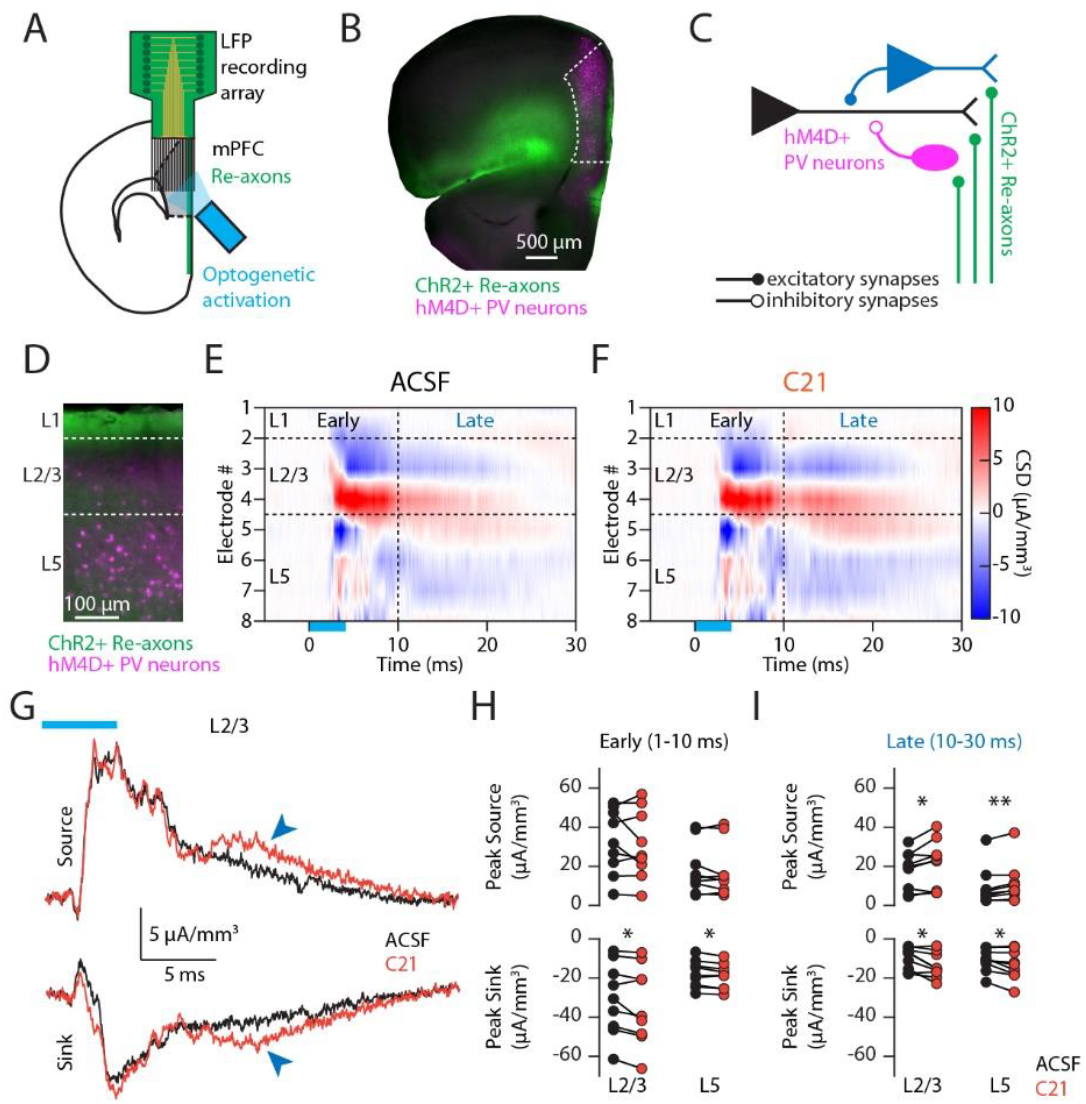
Local field potential recordings in mPFC slices revealed prolonged synaptic activity upon Re optogenetic activation. (A) Experimental approach: whole-cell patch-clamp recording of mPFC neurons combined with optogenetic stimulation of Re axons. (B) Epifluorescent micrograph of a coronal brain slice across mPFC (dashed white line) from a PV-cre mouse. Re axons can be observed in mPFC layer 1 (green). hM4D+ PV neurons (magenta) are spread across mPFC layers. (C) Schematic illustrating the recruitment of excitatory (black and blue) and inhibitory PV positive neurons (magenta) in mPFC by Re axons. (D) Epifluorescent micrograph of a cross section of mPFC. Green: Re axons. Magenta: hM4D+ PV neurons. (E) Color image plot of CSD data from mPFC after optogenetic activation of Reuniens afferents. Layers and electrodes are aligned with the cortical layers shown in D. Cool colors (blue) represent current sinks, and hot colors (red) represent current sources; white is approximately zero. Horizontal dashed lines indicate cortical layers boundaries, as in D. The vertical dashed line at 10 ms after light onset separates early synaptic activity from late synaptic activity. Note that the colors for peak early sinks and sources are saturated to highlight the late responses. (F) Same as in D after bath application of compound 21 (C21), an activator of the hM4D that leads to hyperpolarization of expressing PV neurons. (G) L2/3 CSD traces from (E) and (F) showing the increase in late signals (blue arrow heads) after C21 bath application. (H) Scatter plot of the peak source (top) and peak sink (bottom) resulting from early synaptic activity in L2/3 and L5 mPFC in ACSF (black) and in C21 (orange). (I) Same as F for late synaptic activity. Significance: * p < 0.05, ** p < 0.01, and *** p < 0.001.

To disentangle between feedforward inhibition and excitation, we blocked the inhibitory signal in a cohort of C57BL6/J mice with bath application of gabazine, a GABA_A_ receptor blocker (Suppl. Fig 6AB). If the late (10-30 ms) synaptic activity were dominated by inhibitory signal, one could expect a reduction in CSD peak. However, we observed a strong increase in sinks (L2-ACSF: -7+/-1 µA/mm^3^, L2-GBZ: -18+/-4 µA/mm^3^, Paired Student’s *t* test, p=0.006 ; L5-ACSF: -4+/-1 µA/mm^3^, L5-GBZ: -14+/-2 µA/mm^3^, Wilcoxon Sign Rank test, p=0.0001) and sources (L2-ACSF: 5+/-1 µA/mm^3^, L2-GBZ: 20+/-3 µA/mm^3^, Wilcoxon Sign Rank test, p=0.00006 ; L5-ACSF: 4+/-1 µA/mm^3^, L5-GBZ: 8+/-1 µA/mm^3^, Paired Student’s *t* test, p=0.0001) across layers (Suppl. Fig 6D). These results are consistent with the hypothesis that feedforward/recurrent excitation contributes to the late synaptic activity. However, we also observed epileptiform runaway excitation (23) upon gabazine application with strong delayed sinks and sources that lasted for tens to hundreds of ms (Suppl. Fig 6E).

To avoid epileptiform responses in the slice, we used a complementary disinhibitory approach using DREADDs (24) to unmask polysynaptic excitation. With the slices prepared from PV-Cre mice expressing hM4D in mPFC PV neurons (Fig 3FG), we bath applied compound 21 (C21), a DREADD agonist, resulting in suppression of firing in PV inhibitory cells and therefor at least a partial blockade of inhibitory signaling. C21 had little to no effect on the early responses, but induced a small increase in the early sinks in L2/3 and L5 (L2-ACSF: -31+/-6 µA/mm^3^, L2-C21: -34+/-7 µA/mm^3^, Paired Student’s *t* test, p=0.037 ; L5-ACSF: -18+/-2 µA/mm^3^, L5-C21: -19+/-2 µA/mm^3^, Paired Student’s *t* test, p=0.022) with no change of the early sources (L2-ACSF: 33+/-6 µA/mm^3^, L2-C21: 31+/-6 µA/mm^3^, Paired Student’s *t* test, p=0.47 ; L5-ACSF: - 18+/-4 µA/mm^3^, L5-C21: -18+/-5 µA/mm^3^, Wilcoxon Sign Rank test, p=0.65) (Fig 3H). This is consistent with the idea that early synaptic activity is mostly dominated by monosynaptic excitatory connections. Furthermore, this partial blockade of PV interneuron activity led to an increase in both polysynaptic (10 to 30 ms post-light stimulus window) sinks (L2-ACSF: -10+/-2 µA/mm^3^, L2-C21: -14+/-2 µA/mm^3^, Paired Student’s *t* test, p=0.015 ; L5-ACSF: -11+/-2 µA/mm^3^, L5-C21: -13+/-3 µA/mm^3^, Paired Student’s *t* test, p=0.026) and sources (L2-ACSF: 18+/-3 µA/mm^3^, L2-C21: 21+/-4 µA/mm^3^, Paired Student’s *t* test, p=0.026 ; L5-ACSF: -10+/-3 µA/mm^3^, L5-C21: -11+/-4 µA/mm^3^, Wilcoxon Sign Rank test, p=0.012) in response to light stimulation of ChR2 in Re afferents (Fig 3I). The direct effect of C21 on hM4D expressing PV neurons is likely a reduction of their GABAergic output, which could have resulted in a reduction of the late sinks and sources. However, we observed an increase in the late sinks and sources, reminiscent of the late EPSCs observed in single cell recordings (Fig 2B), strongly suggesting that Re axons recruit local disynaptic feedforward excitation in mPFC. In a later time window, 30 to 400 ms after light stimulation, there was little to no synaptic activity left in mPFC, and no significant effect of C21, confirming that partial blockade of PV neurons did not cause epileptiform activity in the slice.

Electrical stimulation of the layer 1 of mPFC yielded similar results, with polysynaptic (10-30 ms post-stimulus time window) sinks and sources across the cortical column increasing in intensity following bath application of gabazine (Suppl. Fig 7). These results demonstrate that mPFC amplification of Re inputs involves recurrent activity within the cortical column, and that this late excitatory activity competes with concurrent feedforward inhibition. The observed similarity in prolonged mPFC activation following both electrical stimulation of layer 1 and optogenetic stimulation of Re axons strongly implies that these specialized characteristics of the mPFC network, amplifying Re inputs, may extend to other higher-order thalamic inputs targeting mPFC layer 1, such as those from the mediodorsal and ventromedial thalamus.

### Activation of Re leads to prolonged synaptic activity and unit firing in mPFC in vivo

The intracellular and LFP recordings in slices revealed that Re can induce feedforward excitation in mPFC. This feedforward excitation was enhanced by either chemogenetic inhibition of PV cells or blockade of GABA_A_R. To address whether the cortical amplification of thalamic input in mPFC is present in vivo, we recorded LFP and single units in the mPFC of four C57BL6/J mice using Neuropixels probes, combined with optogenetic activation of Re (Fig 4A). Probes were acutely inserted at a 45-degree angle and traversed mPFC in both hemispheres (Fig 4B). Probe tracks were reconstructed in 3D using the SHARP-Track Matlab suite (Fig 4C). Optogenetic activation of Re induced a succession of LFP deflections measured from electrodes located in the mPFC. The CSD derived from these local field potentials confirmed that the source of synaptic activity was present across L2/3 and L5 of mPFC (Fig 4DE). Pairs of sinks and sources were visible within 10 ms after the optogenetic stimulation, likely corresponding to monosynaptic activation of Re-mPFC connections, and at later time points (10 to 30 ms), consistent with polysynaptic recruitment of the local mPFC network and reminiscent of those observed in vitro (Fig 3). These late sink/source pairs are likely the result of Re recruitment of both feedforward excitation and feedforward inhibition and cannot be disentangled with pharmacology as we did in vitro. However, we hypothesized that if Re activation results in both direct and disynaptic recurrent excitation of mPFC, a single Re stimulation will result in prolonged firing and/or bursting of mPFC that would last tens of milliseconds, like the prolonged synaptic activity revealed through CSD analysis (Fig 3 and 4D) and late synaptic currents in single cells (Fig 1&2).

**Figure 4.**
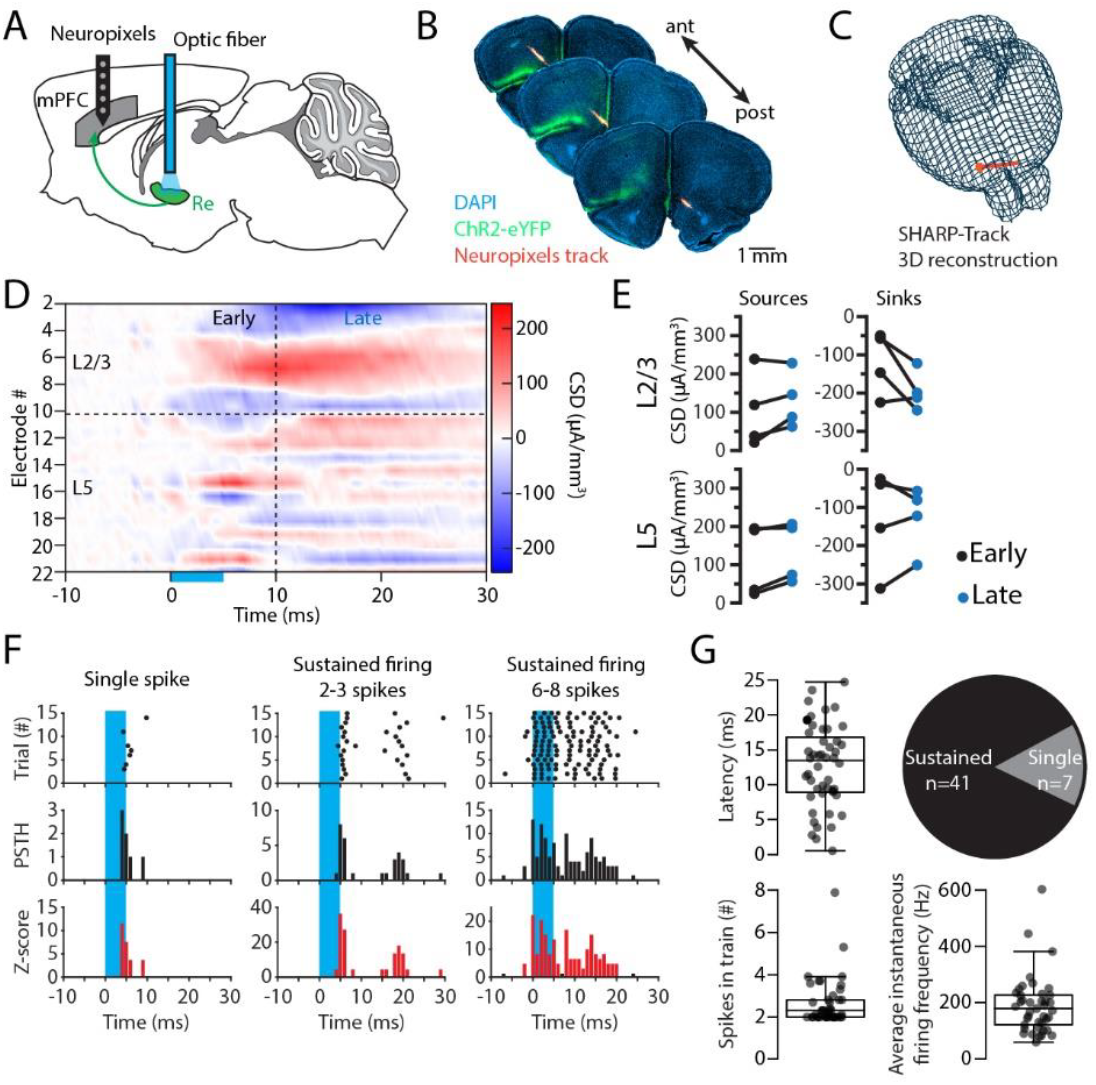
Activation of Re leads to prolonged synaptic activity and unit firing in mPFC in vivo. (A) Scheme showing the Neuropixels recording of mPFC combined with optogenetic stimulation of Re axons. (B) Sequence of epifluorescent micrographs of coronal brain slices through mPFC of a mouse implanted with a Neuropixels probe. The track of the Neuropixels probe is labeled in red. Blue: DAPI staining. Green: ChR2-eYFP axons from the Re. (C) Representation of the Neuropixels probe track through the motor areas and mPFC reconstructed with Allen CCF tools (The Cortical Processing Laboratory at UCL). (D) Color image plot of CSD data from mPFC after optogenetic activation of Reuniens afferents. Cool colors (blue) represent current sinks, and hot colors (red) represent current sources; white is approximately zero. Horizontal dashed lines indicate cortical layers boundaries. The vertical dashed line at 10 ms after light onset separates early synaptic activity from late synaptic activity. (E) Scatter plot of the peak source (left) and peak sink (right) resulting from early synaptic activity (black) and late synaptic activity (blue) across mPFC layers. (F) Raster plots, cumulative histograms, and Z-score analyses from 3 example units responding to a 5 ms blue light activation of Re. (G) Top left: Box-and-whisker plot of the latency from light onset to first spike in mPFC units. Top right: proportion of units responding with single spike (grey) or multiple spikes (black). Bottom: Box-and-whisker plots of the number of spikes in each response train (left) and the average instantaneous spike frequency (right).

Overall, 48 out of 135 isolated mPFC units significantly changed their firing upon activation of Re (Fig 4FG & Suppl. Fig 8). The latency from light onset to initial firing was similar across mPFC layers, with an average of 12.6+/-0.9 ms (n=48) (Fig 4G). Furthermore, 41 out of 48 units responded to Re activation with a sustained firing of 2 to 8 spikes (mean: 2.7+/-0.2 spikes) with a mean instantaneous firing rate of 191+/-16 Hz (Fig 4G). This prolonged firing is consistent with the prolonged synaptic signal observed in LFP/CSD recordings.

Interestingly, beyond this prolonged initial firing, 28/48 responding units showed at least a 50% decrease in their firing 30 to 100 ms after light onset compared to their pre-stimulus firing rate. We also observed a subsequent increase in firing compared to pre-stimulus firing 100 to 400 ms after light onset (Suppl. Fig 8) (Friedman’s test p=2.2×10^−16^, Wilcoxon signed rank tests, Baseline vs Initial p=4.3×10^−13^, Baseline vs Reduction p=0.2, Baseline vs Delay p=1.6×10^−7^). This delayed increase in firing that occurs ∼150 ms after the light onset likely reflect polysynaptic activity, for example Re**→**mPFC**→**mPFC or Re**→**Hippocampus**→**mPFC.

Taken together, the results of these in vivo recordings show that Re recruits long lasting recurrent synaptic activity in mPFC resulting in sustained firing.

## Discussion

The mPFC and hippocampus are well-known for their roles in memory formation and retrieval (1, 25, 26). However, how the interactions between these two structures underlies these functions is not fully understood. The direct pathway between vCA1 and mPFC is involved in social memory and emotional regulation (27). However, dCA1, a region critical for spatial and contextual memory (28), show little to no direct projections to mPFC (29, 30). Similarly, mPFC show little to no direct projections to hippocampus (25). The mPFC plays a role in directing the retrieval of contextual memories even in the absence of a direct pathway to the hippocampus (25), revealing the importance of indirect pathways, notably through the Re in the midline thalamus, for spatial and emotional memory processing.

The Re sends and receives inputs to and from mPFC and hippocampus and can modulate the activity of both structures (3). Our data showed that Re sends monosynaptic excitatory projections to mPFC neurons in L1, L2/3 and L5. Re-mPFC synapses showed a short-term depression profile, supporting the idea that they are high release probability synapses (31). This short-term depression profile at 10 Hz is consistent with a previous report using similar whole-cell patch-clamp and optogenetic approach in rats (20). Optogenetic activation of Re-mPFC synapses also resulted in strong feedforward inhibition of pyramidal cells, consistent with previous reports (32–34). Interestingly, despite this, Re activation produced a strong polysynaptic feedforward excitation of mPFC pyramidal cells, which was particularly evident upon partial blockade of the feedforward inhibition. Activation of Re-EC or PO-S1 synapses did not result in a similar feedforward excitation, suggesting a specialized amplifying feature of the Re-mPFC subsystem. Consistent with the observations of late EPSCs in single cell recordings and prolonged synaptic activity in transcolumnar CSD recording in slices, single unit *in vivo* showed a similar prolonged firing of mPFC network upon activation of Re inputs.

These results suggest that the indirect pathway between hippocampus and mPFC via the Re can recruit recurrent and long-lasting activity in mPFC. This activity could act as a gating mechanisms or coincidence detector for direct input from vCA1 to mPFC, contributing to hippocampal-mPFC synchrony for contextual cue encoding (35).

The prolonged excitation of the mPFC upon activation of Re afferents is likely mediated by direct excitation combined with polysynaptic feedforward excitation. Cortical columns contain major interactions between excitatory cells across layers, which is preserved across the neocortex and plays a major role in the integrative function of the cortex (36). Local connectivity between L2/3 and L5 of mPFC is a likely candidate to explain the prolonged excitation of mPFC upon activation of Re. Both L2/3 and L5 pyramidal cells receive direct synaptic inputs from Re (3), which can lead to action potential discharge from neurons in both layers. Recurrent connectivity within and across layers in mPFC (37, 38) could then lead to a second, late excitation. Interestingly, higher order thalamic nuclei have been shown to facilitate generation of dendritic plateau potentials in cortical pyramidal neurons (39). Re axons in L1 likely target apical dendrites of L2/3 and L5 pyramidal cells. Glutamate releases onto synapses in these apical dendrites could generate dendritic plateau potentials, putting the neuron in an “activated state”. This activated state has been shown to operate as a time window on the order of hundreds of ms during which cortical neurons are particularly excitable and more likely to respond to afferent inputs with spike discharge (40). This phenomenon could create a temporal window during which other inputs to mPFC could be amplified, notably the direct pathway from vCA1 that is important for contextual cues encoding.

It is important to note that the small size and position of Re makes it challenging to specifically target it with viral injection, and this is a potential limitation of our approach. Some spillover of viral vectors into neighboring midline and intralaminar thalamic nuclei that sends projections to mPFC can lead to expression of ChR2 in their axons (41). Stimulation of the afferents from these multiple thalamic nuclei could contribute to the sustained excitation of mPFC neurons. Similarly, polysynaptic connections from Re to other cortical/subcortical structures, then to mPFC could explain the prolonged excitation. While this possibility cannot be excluded during the *in vivo* recordings (Fig. 4) where the brain is fully intact, this is unlikely in ex vivo slice recording as only local axons are intact (Fig. 1&3).

Electrical stimulation of L1 produced a similar prolonged excitation in the mPFC columns than optogenetic activation of Re afferents (Suppl. Fig. 7), suggesting that a) much of the Re activation of mPFC arrives through this layer, and b) activation of other structures projecting to mPFC through L1, such as the mediodorsal thalamus, would result in a similar amplification. Interestingly, this amplification of thalamic input seems to be uniquely prominent in mPFC. Single cell recording in EC, another cortical target of Re, did not reveal late EPSCs. We also recorded a subset of somatosensory cortex pyramidal neurons while optogenetically stimulating the posterior thalamus, another higher order thalamic structure. Only 2/10 cells responded with a late EPSCs, compared to the 50% in mPFC upon Re activation. These results further suggest that the mechanism underlying prolonged excitation resides in the local network connectivity in mPFC rather than the Re outputs.

The mPFC plays a crucial role in integrating information from its numerous cortical and subcortical afferents and to converge updated information to its targets (42). The prolonged excitation of mPFC neurons we observed is significant as it can amplify the transmission of information to its targets. The propensity of the mPFC for prolonged excitation may act as an integrator where a burst discharge occurs upon synchronized inputs to mPFC. Overall, our work suggests that mPFC, but not EC, contains the necessary local circuitry for sustaining Re coordinated information transferred between these memory and executive function related brain regions.

This study provides robust physiological characterization of the synaptic properties of the Re-mPFC and reveals an unprecedented recruitment of feedforward excitation of the mPFC by a thalamic input. This prolonged excitation may underly the role of mPFC in integrating information, maintain it in working memory, and in providing a strong overriding output to its postsynaptic targets upon synchronized input to mPFC.

### Materials and Methods

### Experimental model and subject details

All experimental procedures were performed in accordance with the guidelines of the Institutional Animal Care and Use Committees of Stanford University. Young adult (4 to 8 weeks old) C57BL/6J (M. musculus, Jackson Laboratory, RRID: IMSR_JAX:000664) PV-Cre (M. musculus, Jackson Laboratory, RRID: IMSR_JAX:008069) mice of either sex were housed prior to experiment in a temperature and humidity-controlled animal house. They were maintained in a 12 h/12 h light-dark cycle and could access food and water ad libitum.

### Viral injections

The procedure for viral injection has been described in detail in (24). Briefly, young adult mice were anesthetized with Ketamine-Xylazine (83 and 3.5 mg/kg, respectively) or isoflurane (5% induction, 2% maintenance) and received Carprofen 5 mg/kg i.p. Mice received injections in Re (in mm from bregma, AP -0.75, ML 0, DV -4), mPFC (AP +2, ML +/-0.2, DV -1.7) or PO (AP -1.7, ML +/-1.25, DV -2.9). A volume of 0.2 to 0.5 µl of virus encoding the ChR2 (AAV1-CaMKIIa-ChR2(H134R)_eYFP-WPRE-HGH, 10^12^ GC, 100 nl/min, Penn Vector Core, Addgene 26969P) was delivered at the injection site in Re or PO. A volume of 0.5 µl of virus encoding the inhibitory DREADD (AAV8-hSyn-DIO-hM4D(Gi)-mCherry, 10^13^ GC, 100 nl/min, Penn Vector Core, Addgene 44362-AAV8) was delivered in mPFC.

### In vitro patch-clamp recording

#### Solutions for slice preparation and recording

Mouse brains were extracted, sliced with a vibratome (Leica VT1200S) and recorded using standard procedures (24). In brief, 300 µm-thick coronal brain slices were obtained in an ice-cold oxygenated slicing solution containing in mM: 66 NaCl, 2.5 KCl, 1.25 NaH_2_PO_4_, 26 NaHCO_3_, 105 D(+)-sucrose, 27 D(+)-glucose, 1.7 L(+)-ascorbic acid, 0.5 CaCl_2_ and 7 MgCl_2_. Slices were then kept in a recovery chamber at 35°C for 30 min then at room temperature for at least 30 more minutes before recording. The chamber was filled with a recovery solution containing, in mM: 131 NaCl, 2.5 KCl, 1.25 NaH_2_PO_4_, 26 NaHCO_3_, 20 D(+)-glucose, 1.7 L(+)-ascorbic acid, 2 CaCl_2_, 1.2 MgCl_2_, 3 myo-inositol, 2 pyruvate. The artificial cerebrospinal fluid (ACSF) used for recording contained, in mM: 131 NaCl, 2.5 KCl, 1.25 NaH_2_PO_4_, 26 NaHCO_3_, 20 D(+)-glucose, 1.7 L(+)-ascorbic acid, 2 CaCl_2_, 1.2 MgCl_2_, 0.01 glycine and was supplemented with 0.1 picrotoxin (Abcam, ab120315) when appropriate. Cesium-based intracellular solution contained in mM: 127 CsGluconate, 10 HEPES, 2 CsBAPTA, 6 MgCl_2_, 10 phosphocreatine, 2 Mg-ATP, 0.4 Na-GTP, 2 QX314-Cl (Tocris, 2313), pH 7.3, 290–305 mOsm. The potassium-based intracellular solution contained in mM: 144 KGluconate, 10 HEPES, 3 MgCl_2_, 0.5 EGTA, pH 7.3, 290–305 mOsm. Series and input resistances were monitored throughout recordings by brief voltage and current pulses, for voltage- and current-clamp recordings, respectively. Data were excluded for Rs and Ri changes above 20%. Recordings are not corrected for a liquid junction potential of -10 mV.

#### Properties of excitatory synapses

ChR2-expressing afferents were activated using blue LEDs stimulation (Cairn Res, Faversham, UK, 455 nm, duration: 1 ms, maximal light intensity 3.5 mW, 0.16 mW/mm^2^; Thorlabs M450LP2, 450 nm, duration 1 ms, maximal light intensity 13 mW, 19 mW/cm^2^) or blue lasers (Laserglow Technologies, 473 nm, duration 1 ms, maximal light intensity 8 mW, 10.8 mW/cm^2^).

To confirm the monosynaptic nature of Re-mPFC connections, single light pulses were used to activate Re afferents once every 10 s. Action potential-dependent light evoked EPSCs were blocked with tetrodotoxin (TTX, Latoxan L8503) 0.25 µM. Action potential-independent light evoked responses were then recorded subsequent to bath application of 4-aminopyridine (4AP, Abcam, ab120122) 100 µM.

The subunit content of the Re-mPFC synapses was assessed using successive bath application of ionotropic glutamate receptor blockers in the presence of picrotoxin (Sigma-Aldrich, P1675), a GABA_A_ receptor blocker. The AMPA receptor mediated currents were recorded at -70 mV for at least 2 min of stable baseline and then blocked with 6,7-dinitroquinoxaline-2,3(1H,4H)-dione (DNQX, Abcam, ab120169) 40 µM. The membrane potential was brought at +40 mV to release the magnesium block and show NMDA receptor mediated currents. After a baseline recording, the NMDA receptors were partially or completely blocked using NVP-AAM077 (NVP, Novartis Pharma) 0.05 µM (a competitive antagonist with higher selectivity for GluN2A containing NMDA receptors (43)), CP-101.606 (CP, Pfizer Pharmaceuticals) 10 µM (GluN2B specific blocker (44)), (2S*,3R*)-1-(Phenanthren-2-carbonyl)piperazine-2,3-dicarboxylic acid (PPDA, Tocris Bioscience, 2530) 0.5 µM (NMDA receptor antagonist that preferentially binds to GluN2C/GluN2D (45)) or DL-2-Amino-5-phosphonovaleric acid (APV, Sigma, A5282) 100 µM (broad spectrum NMDA receptor blocker).

To arrive at a simple index of synaptic response duration, we calculate the weighted time constant (τ_w_), as by dividing the total synaptic charge (area under the curve of the composite EPSC) by the peak EPSC amplitude. The unit of τ_w_ is time (ms) and provides an approximation of the amount it takes to deliver a fraction (1/*e*) of the total synaptic charge. Larger values of τ_w_ would correspond to polysynaptic composite responses, especially compared to isolated monosynaptic responses.

#### Feedforward inhibition

mPFC and EC neurons were recorded in voltage-clamp at -70 mV then at +10 mV to show EPSC and IPSC respectively. The IPSCs were then blocked with Gabazine (Abcam, ab120042) 10 µM in a subset of experiments.

#### Cellular properties and synaptic depolarization

Passive cellular properties were recorded in voltage clamp at resting levels of -60 to -70 mV, using a 500 ms-long hyperpolarizing step of 10 mV or in current-clamp from the resting membrane potential. Action potential properties were measured at the rheobase in current-clamp mode, with a membrane potential held at ∼ -60 mV and using 500 ms-long depolarizing current steps of increasing amplitude.

### In vitro local field potential recordings

Extracellular recordings were conducted using 400 µm-thick coronal brain slices containing the mPFC obtained from PV-Cre and C57BL6/J mice expressing the ChR2-eYFP in Re after viral injection. The experimental setup involved placing the slices in a humidified, oxygenated interface chamber at a constant temperature of 34°C and perfusing them with oxygenated ACSF at a rate of 2 ml/min. Local field potentials (LFPs) were measured with a linear silicon multichannel probe (NeuroNexus Technologies) with 16 channels and 100 µm inter-electrode spacing, which was placed perpendicular to the laminar plane across all mPFC layers.

The multichannel probe was connected to a pre-amplifier (PZ5-32, Tucker-Davis Technologies (TDT)), and data were stored using a RZ5D processor multichannel workstation (TDT). Recording sessions were designed with the Real-time Processor Visual Design Studio (RPvdsEx) tool and custom Python code. LFP data were acquired at 25 kHz frequency and band-passed between 0.1 and 500 Hz.

Optogenetic stimulations were delivered using a blue laser (Laserglow Technologies, 473 nm, duration 5 ms, maximal light intensity 15 mW, 20 mW/cm^2^). The blue light stimulation encompassed a circular area of about 1 mm in diameter spanning across all cortical layer in mPFC. Electrical stimulations were delivered by a bipolar tungsten microelectrode (with each wire having a resistance of 50-100 kΩ, duration 0.1 ms, intensity 500 µA) that was precisely positioned in layer 1 of the mPFC. Both stimulations were done as a single pulse, once every 20 s. Each successive recording conditions (ACSF, C21 or Gabazine, DNQX/AP5, TTX) were recorded for 5-8 min but only the last 2 min were used for analysis, leaving enough time to reach a steady state in the different drug conditions.

To accurately characterize the location, direction, and magnitude of currents associated with the synchronous network activity, we derived the current-source density (CSD) from the raw LFP signals (21). We assumed a uniform conductivity of 0.3 S/m across all layers (46). The CSD was computed in the mPFC by performing second order algebraic differentiation of the LFP signals recorded at adjacent electrodes in the linear probe. The ion flow in and out of the extracellular space created sinks and sources of current in the CSD, which can then be attributed to specific cortical layers. Successive pharmacological blockade of postsynaptic receptors and action potential generation allowed to disentangle the origins of the recorded sinks and sources in mPFC.

### Acute *in vivo* Neuropixels recordings

#### Mouse preparation

Four female C57BL6/J mice (P110-P120) were injected with AAV1-CaMKIIa-ChR2(H134R)_eYFP-WPRE-HGH as described above. An optical fiber (200 μm core, 0.22-NA, 2.5 mm-diameter ceramic ferrule, Thorlabs) was implanted 200 µm above the viral injection site. A miniature self-tapping screw (stainless steel, J.I Morris Company FF00CE125) connected to a pin (Mill-Max 853-93-100-10-001000) was inserted into the occipital bone over the cerebellum as a reference. A gold pin (1.27 mm, Mill-Max Manufacturing Corp.) was implanted in contact with the surface of primary somatosensory cortex and served as an EEG electrode to validate that no pathological activity, e.g., seizures, were induced. A custom metal head bar (eMachineShop) was placed and stabilized using dental cement (Metabond). After surgery, animals recovered for one week before being trained for handling and head restriction on a cylindrical treadmill. Recordings were performed after habituation over one to two weeks of training.

On recording day 1, a 1 mm craniotomy was performed with a 0.5 mm burr dental drill (Fine Science Tools) over the right motor cortex (AP: 1.5 to 2.5 mm, ML: 1 to 2 mm) and sealed with a removable silicone sealant (Kwik-Cast, World Precision Instrument). Mice were left to recover for at least 4h in their home cage before recording. Mice were head fixed on the cylindrical treadmill and an optic fiber was connected via the ceramic ferrule/optic fiber previously mounted on the mouse. The other end of the optic fiber was connected to a laser (OEM Laser Systems, 473 nm, duration 5ms, maximal light intensity 3.2 mW, 4.5 mW/cm^2^), while the EEG gold pin and reference screw were connected to an Intan headstage (RHD 2132) and OpenEphys acquisition board. The silicone sealant was removed from the craniotomy and a Neuropixels 1.0 probe previously dipped in red dye (Vybrant DiD cell labeling solution, ThermoFisher Scientific) was connected via cable to the PXIe data acquisition system (National Instrument NI-PXIe-1071) then positioned above the cortex.

#### Recording

The Neuropixels probe was slowly inserted over ∼5 minutes using a Sutter MP285 micromanipulator system at a 45-degree angle into the brain of the awake, head-fixed mouse, followed by about an hour of recording. This electrode array recorded 384 channels, spanning 3.8 mm of the brain across motor cortex and mPFC, traversing the midline. The probe was allowed to settle for ∼20 min before starting the recording. Recording sessions consisted of 5 min of baseline followed by 5 min of optogenetic stimulation at either 1 or 10 Hz. Trains of stimulation were given once every 20 s and consisted of 2 (at 1 Hz) or 11 (at 10 Hz) 5-ms long pulses. Data were acquire using OpenEphys software (https://open-ephys.org/) at 30 kHz.

#### Analysis

The data were spike-sorted automatically by Kilosort2.5 (https://github.com/MouseLand/Kilosort) followed by manual curating using Phy2 (https://github.com/cortex-lab/phy). Single unit clusters with inter-spike-interval violations > 0.05% (calculated as the ratio of the spikes within the refractory period ±1.5 ms to the total number of spikes) were removed. Single unit clusters that were not stable throughout the 10 min recording sessions were excluded from analysis. Custom Matlab code (www.mathworks.com) were used to plot peri-stimulus raster plots and peri-event time histogram (in 1 or 5 ms bin) from 200 ms before to 400 ms after laser stimulation. Z-scores were calculated using the 200 ms prior to laser stimulation as a baseline.

#### Histology and probe localization

Mice were sacrificed with an intraperitoneal injection of Fatal+ (Pentobarbital, Vortech Pharmaceutical). Cardiac perfusion was performed with 4% paraformaldehyde in PBS. The brains were post-fixed overnight in 4% paraformaldehyde, rinsed in PBS and sliced using a vibratome (Leica VT1000S). The 100 µm-thick brain slices containing the recording site in mPFC and the viral injection site in the Re were mounted in DAPI-Fluoromount-G (Southern Biotech 0100-20). Epifluorescent micrographs for DAPI, ChR2-eYFP and probe track were obtained using a Zeiss microscope (Zeiss Axio Imager M2). SHARP-track (Allen CCF tools) was used to reconstruct the Neuropixels probe track in 3D and to localize recording sites. SHARP-track allows the 3D reconstruction of the Neuropixels probe within the Allen Mouse Brain Common Coordinate Framework based on the histology and physiological landmarks. Each channel was then attributed to its corresponding brain regions in motor cortex or mPFC. The mPFC includes orbital areas, prelimbic cortex, infralimbic cortex, cingulate cortex, frontal pole (or area 24, area 25 and area 32 from Paxinos 5th edition).

### Quantification and statistical analysis

The R programming software (3.6.1, The R Foundation for Statistical Computing, 2019] and Prism 6 (GraphPad Software Inc) were used for statistical analysis. The Shapiro-Wilk normality test was used to assess the normality of the dataset distribution. Accordingly, comparison of two datasets were done using Student’s *t* test and paired Student’s *t* test for non-repeated and repeated measures, respectively. Their non-parametric equivalent, Wilcoxon rank sum test and Wilcoxon signed rank test, were used when at least one of the datasets was not normally distributed. Comparisons of more than 2 datasets were done using ANOVA tests or their non-parametric equivalents when appropriate. The Šidák correction was used to adjust the α threshold for significance when the analysis required multiple comparisons. Significant exact p values are indicated in the main text. On figures, significance levels are abbreviated as * (p < 0.05), ** (p < 0.01) and *** (p < 0.001). Detailed statistics can be found in Table 1.

## Acknowledgments

We thank Prof. Anita Lüthi for providing lab space, equipment, and overall support during the initial phase of the project. We thank all lab members for constructive input for the manuscript and discussions during the project. We thank Nicole Agranonik, Cameron Glick and Katlin Villar for their help in managing mouse lines, laboratory equipment and supplies.

