## Supplemental Figures and Tables for "Reuniens thalamus recruits recurrent excitation in medial prefrontal cortex"

**Supporting Information for**
### Reuniens thalamus recruits recurrent excitation in medial prefrontal cortex.

Gil Vantomme, Gabrielle Devienne, Jacob M Hull, John R Huguenard*

John R Huguenard


**This PDF file includes:**

Figures S1 to S8
Tables S1 to S2

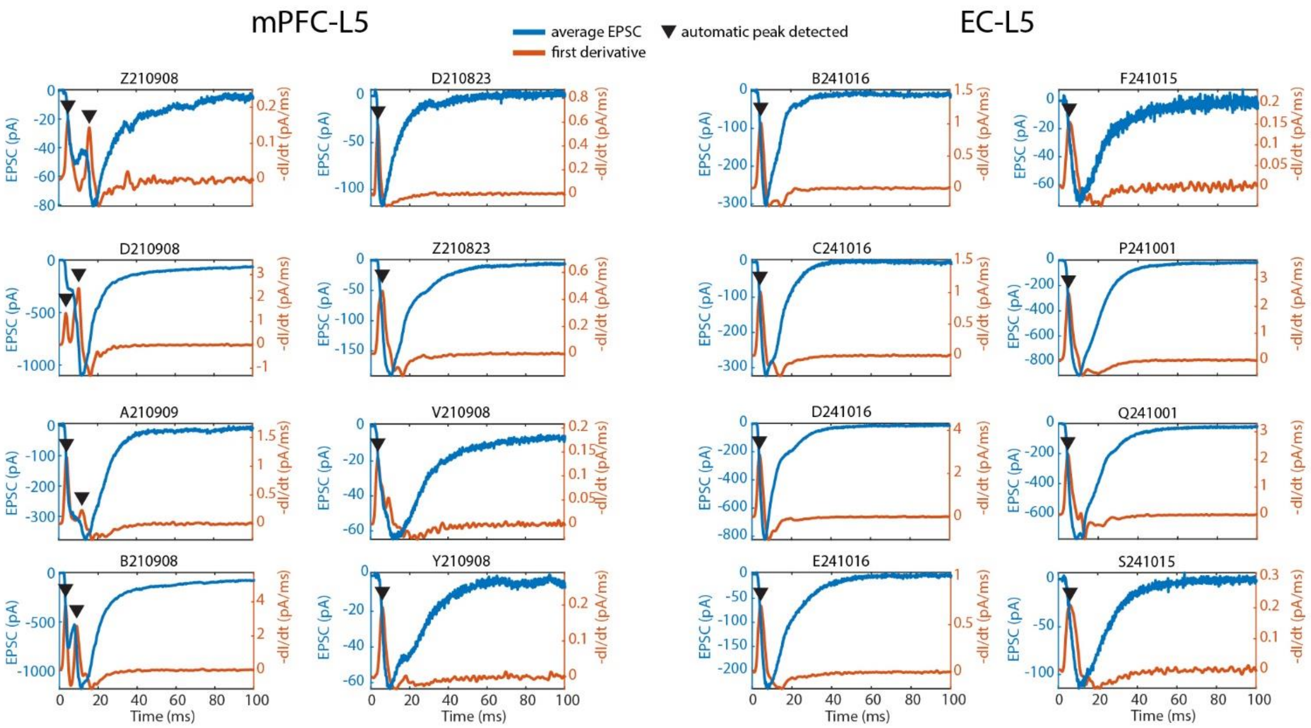


**Fig. S1. Examples of automated EPSC peak detection.** Average EPSCs (blue) were derived once over time (orange). Peaks on the first derivative were detected if the rate of change and the minimum peak prominence were at least 0.1 pA/ms (black arrowheads).

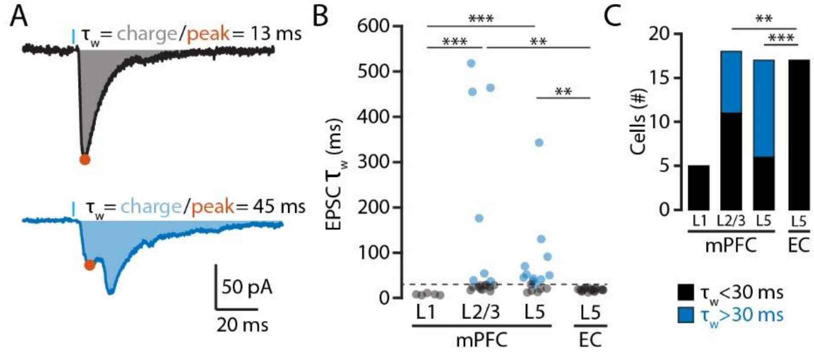


**Fig. S2. EPSCs in L2/3 and L5 mPFC have longer weighted decay time constant ($\tau_w$) than in EC.** (A) Example traces of EPSCs showing how $\tau_w$ was calculated by dividing the charge (area under the curve) by the amplitude of the first peak (orange dot). (B) Scatter plot of the EPSC $\tau_w$ across mPFC layers and in layer 5 EC. (C) Histogram of the number of cells with EPSC $\tau_w$ below (black) and above (blue) 30 ms. Note that the proportion is similar to the number of cells showing a prominent late EPSC in Fig 1E. Significance: * $p < 0.05$, ** $p < 0.01$, and *** $p < 0.001$.

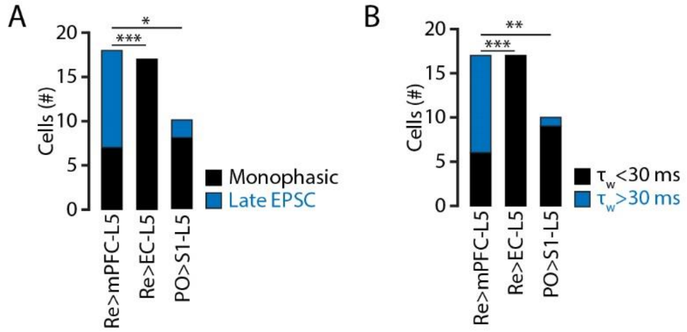


**Fig. S3. Reuniens activation recruits more polysynaptic excitation in mPFC than in EC and more than PO thalamus-somatosensory cortex. (**A) Histogram showing the number of cells with only monophasic EPSCs (black) and cells with prominent late EPSCs (blue). (B) Histogram showing the number of cells with EPSC $\tau_w$ below 30 ms (black) and above 30 ms (blue).

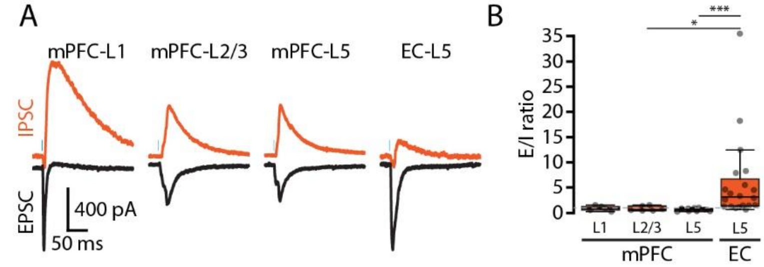


**Fig. S4. Reuniens recruitment of polysynaptic inhibition is greater in mPFC than in EC.** (A) Typical traces of EPSCs recorded at -70 mV (black) and of IPSCs recorded at +10 mV (orange) in mPFC and EC. (B) Excitation/inhibition ratio showing the reduction in recruitment of feedforward inhibition in EC compared to mPFC.

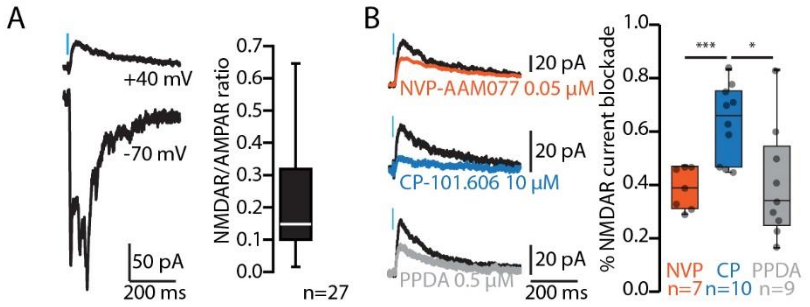


**Fig. S5. Reuniens activation in mPFC recruits NMDARs.** (A) Left, example traces of the AMPAR mediated current recorded at -70 mV in ACSF with picrotoxin (bottom) and the NMDAR mediated current recorded at +40 mV in ACSF, picrotoxin and DNQX (top). Right, NMDAR/AMPAR ratios. (B) Left, representative traces of NMDAR mediated currents recorded in baseline conditions (black) and after NVP (orange), CP (dark blue) or PPDA (grey) bath application. Right, corresponding population data.

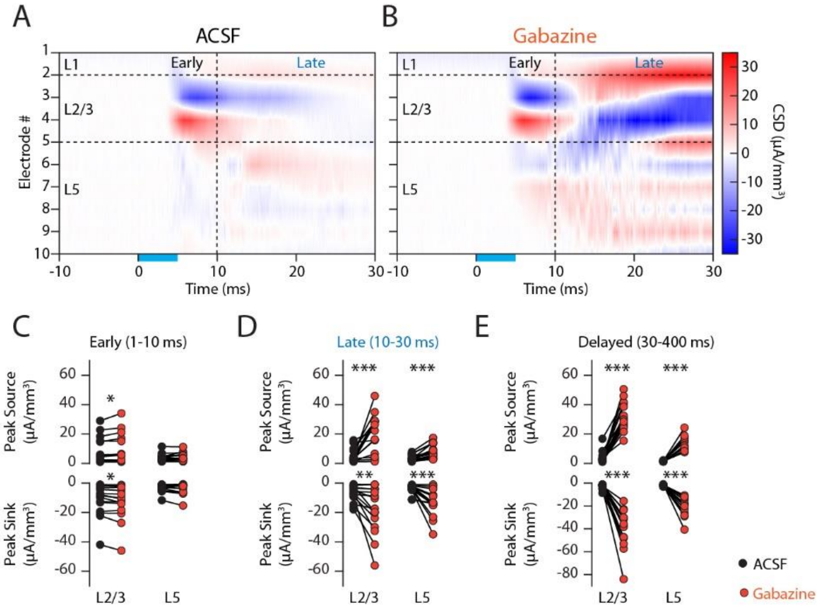


**Fig. S6. GABA$_A$R blockade reveals the delayed activation of mPFC local network upon Re activation.** (A) Color image plot of CSD data from mPFC after optogenetic activation of Reuniens afferents. Cool colors (blue) represent current sinks, and hot colors (red) represent current sources; white is approximately zero. Horizontal dashed lines indicate cortical layers boundaries. Vertical dashed line at 10 ms after light onset separates early synaptic activity from late synaptic activity. (B) Same as in A after bath application of the GABA$_A$R blocker, gabazine. Note the increase in late synaptic activity. (C) Scatter plot of the peak source (top) and peak sink (bottom) resulting from early synaptic activity in L2/3 and L5 mPFC in ACSF (black) and in Gabazine (orange). (D) Same as C for late synaptic activity. (E) Same as (C) for delayed synaptic activity.

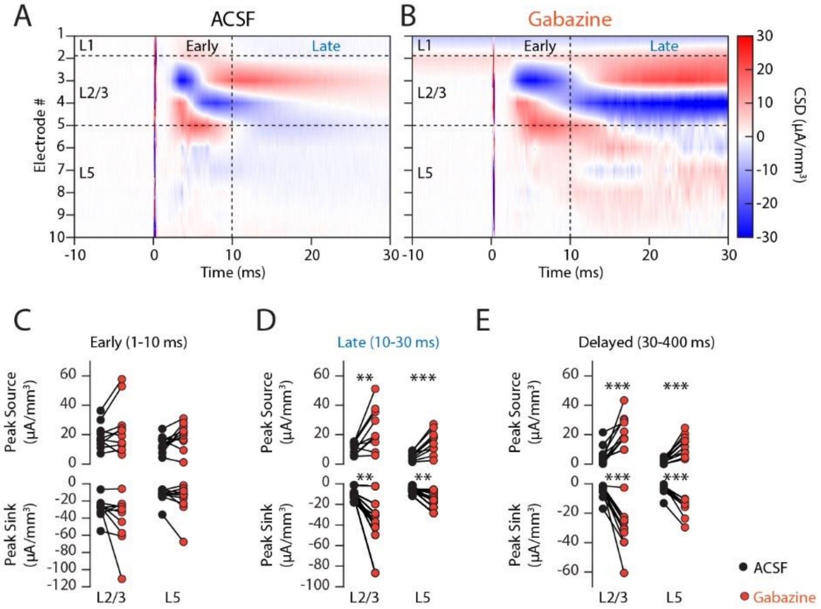


**Fig. S7. Layer 1 stimulation recruits a feedforward excitatory circuit in mPFC in mice.** (A) Color image plot of CSD data from mPFC after electrical stimulation of layer 1. Cool colors (blue) represent current sinks, and hot colors (red) represent current sources; white is approximately zero. Horizontal dashed lines indicate cortical layers boundaries. Vertical dashed line at 10 ms after stimulation onset separates early synaptic activity from late synaptic activity. (B) Same as in A after bath application of the $GABA_AR$ blocker, gabazine. Note the increase in late synaptic activity. (C) Scatter plot of the peak source (top) and peak sink (bottom) resulting from early synaptic activity in L2/3 and L5 mPFC in ACSF (black) and in Gabazine (orange). (D) Same as C for late synaptic activity. (E) Same as (C) for delayed synaptic activity.

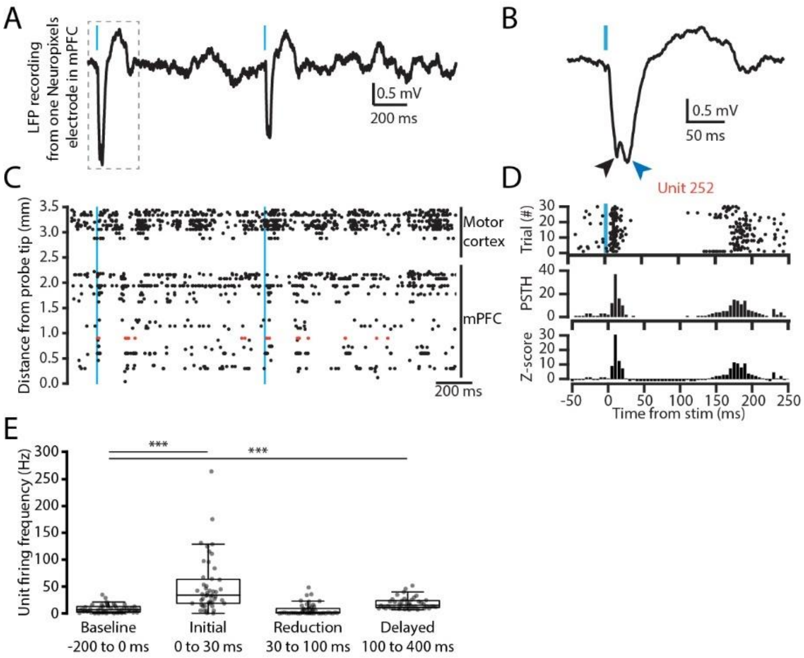


**Fig. S8. Activation of Re leads to biphasic excitation of mPFC units in vivo.** (A) Example trace of local field potential recording in mPFC upon optogenetic stimulation of Re. (B) Expanded trace from D. (C) Raster of 94 units firing during a 2 s long recording. Units are organized by their relative position from the probe tip. Vertical blue lines indicate two stimulations of the Re nucleus. (D) Raster plot, cumulative histogram, and Z-score analysis from an example unit responding to a 5 ms blue light activation of Re. (E) Box-and-whisker plots of the average firing frequency of mPFC units before and after light activation of Re. Significance: * $p < 0.05$, ** $p < 0.01$, and *** $p < 0.001$.

**Table S1. Cellular and EPSC properties.**

| Parameters | mPFC-L1 | mPFC-L2 | mPFC-L5 | EC-L5 |
|---|---|---|---|---|
| Resting membrane Potential (mV) | -65 ± 2, n=7 | -61 ± 3, n=15 | -61 ± 2, n=13 | -63 ± 1, n=19 |
| Cell Resistance (MOhm) | 369 ± 15, n=5 | 405 ± 39, n=15 | 396 ± 42, n=23 | 289 ± 23, n=20 |
| Cell Capacitance (pF) | 35 ± 2, n=5 | 104 ± 8, n=15 | 101 ± 9, n=23 | 149 ± 8, n=20 |
| Rheobase (pA) | 54 ± 11, n=7 | 23 ± 5, n=14 | 39 ± 7, n=13 | 32 ± 3, n=19 |
| AP Threshold (mV) | -34 ± 1, n=7 | -36 ± 1, n=14 | -41 ± 1, n=13 | -38 ± 1, n=19 |
| AP Overshoot (mV) | 20 ± 1, n=7 | 47 ± 3, n=14 | 36 ± 4, n=13 | 47 ± 1, n=19 |
| AP AHP (mV) | -19 ± 2, n=7 | -13 ± 1, n=14 | -12 ± 1, n=13 | -16 ± 1, n=19 |
| AP Half-Width (ms) | 2.2 ± 0.1, n=7 | 2.1 ± 0.1, n=14 | 1.9 ± 0.1, n=13 | 2.2 ± 0.1, n=19 |
| Avg AP Freq at Twice Rheobase (Hz) | 22 ± 3, n=6 | 8 ± 1, n=12 | 13 ± 2, n=12 | 7 ± 1, n=18 |
| EPSC Amplitude (pA) | 278 ± 159, n=6 | 248 ± 42, n=22 | 282 ± 77, n=23 | 149 ± 41, n=20 |
| EPSC Latency (ms) | 2.8 ± 1, n=6 | 4.3 ± 0.9, n=22 | 3.5 ± 0.6, n=23 | 3.2 ± 0.3, n=20 |
| EPSC Rise Time (ms) | 1.3 ± 0.4, n=6 | 3.8 ± 0.6, n=22 | 3 ± 0.5, n=23 | 2.9 ± 0.3, n=20 |
| EPSC Decay Time (ms) | 12 ± 3, n=5 | 22 ± 2, n=13 | 58 ± 21, n=20 | 17 ± 2, n=20 |
| PPR 10 Hz, second/first EPSC | 3.37 ± 2.33, N=6 | 0.9 ± 0.08, N=15 | 1.05 ± 0.11, N=11 | 0.79 ± 0.03, N=20 |
| PPR 10 Hz, tenth/first EPSC | 1.69 ± 1.09, N=6 | 0.32 ± 0.02, N=15 | 0.42 ± 0.07, N=11 | 0.37 ± 0.04, N=20 |

**Table S2. Summary statistics.**

| Fig. panel | Main statistic | post hoc tests | Exact p value |
|---|---|---|---|
| 1E | Chi-square tests | na | mPFC-L5 vs EC-L5, p=0.0001<br>mPFC-L2/3 vs EC-L5, p=0.005<br>mPFC-L1 vs EC-L5, p=0.06 |
| 1F | Wilcoxon rank sum test | na | Latency, $p=9x10^{-10}$ |
| 1H | Friedman rank sum test | Wilcoxon signed rank tests, Šidák correction, α=0.017 | Friedman: p=0.001<br>ACSF-TTX: p=0.016<br>TTX-4AP: p=0.016<br>BL-4AP: p=0.016 |
| Suppl. 2B | Kruskal–Wallis H test | Dunn's Multiple Comparison Test, p-values adjusted with Hochberg method | Kruskal–Wallis H test: $p=5.3x10^{-6}$<br>mPFC-L5 vs EC-L5, p=0.0051<br>mPFC-L2/3 vs EC-L5, p=0.0039<br>mPFC-L1 vs EC-L5, p=0.1<br>mPFC-L5 vs mPFC-L2/3, p=0.96<br>mPFC-L5 vs mPFC-L1, p=0.0003<br>mPFC-L2/3 vs mPFC-L1, p=0.0002 |
| Suppl. 2C | Chi-square tests | na | mPFC-L5 vs EC-L5, p=0.00006<br>mPFC-L2/3 vs EC-L5, p=0.004<br>mPFC-L1 vs EC-L5, NA |
| Suppl. 3A | Chi-square tests | na | mPFC-L5 vs EC-L5, p=0.0001<br>mPFC-L5 vs S1-L5, p=0.037<br>EC-L5 vs S1-L5, p=0.06 |
| Suppl. 3B | Chi-square tests | na | mPFC-L5 vs EC-L5, p=0.00006<br>mPFC-L5 vs S1-L5, p=0.0057<br>EC-L5 vs S1-L5, p=0.18 |
| 2C | Paired Student's *t* test | na | IPSC amplitude, p=0.002 |
| 2D | Wilcoxon signed rank test | na | Late EPSC area, p=0.03 |
| 2F | Friedman rank sum test | Wilcoxon signed rank tests, Šidák correction, α=0.017 | **Late EPSC,**<br>Friedman: $p=3.4x10^{-4}$<br>ACSF vs APV: p= 0.008<br>APV vs DNQX: p=0.008<br>ACSF vs DNQX: p=0.008<br>**Early EPSC,**<br>Friedman: $p=8.1x10^{-4}$<br>ACSF vs APV: p= 0.11<br>APV vs DNQX: p=0.008<br>ACSF vs DNQX: p=0.008 |
| 2C | Friedman rank sum test | Wilcoxon signed rank tests, Šidák correction, α=0.017 | Friedman: $p=3.4x10^{-4}$<br>BL-TTX: p=0.004<br>TTX-4AP: p=0.008<br>BL-4AP: p=0.008 |
| Suppl. 4B | Kruskal–Wallis H test | Dunn's Multiple Comparison Test, p-values adjusted with Hochberg method | Kruskal–Wallis H test: $p=5.9x10^{-5}$<br>mPFC-L5 vs EC-L5, p=0.0001<br>mPFC-L2/3 vs EC-L5, p=0.0276<br>mPFC-L1 vs EC-L5, p=0.052<br>mPFC-L5 vs mPFC-L2/3, p=0.99<br>mPFC-L5 vs mPFC-L1, p=1<br>mPFC-L2/3 vs mPFC-L1, p=0.94 |
| Suppl. 5B | One-way ANOVA | Student's *t* tests, Šidák correction, α=0.017 | One-way ANOVA: p=0.0037<br>NVP vs CP: p=0.0004<br>CP vs PPDA: p=0.01<br>NVP vs PPDA: p=0.83 |
| 3F | Paired Student's *t* test or Wilcoxon signed rank test | na | **C21 effect on early:**<br>L2/3 sink p=0.037 |

| | | | |
|---|---|---|---|
| | | | L5 sink p=0.022<br>L2/3 source p=0.47<br>L5 source p=0.65 |
| 3G | Paired Student's *t* test or Wilcoxon signed rank test | na | **C21 effect on late:**<br>L2/3 sink p=0.015<br>L5 sink p=0.026<br>L2/3 source p=0.026<br>L5 source p=0.012 |
| Suppl. 6C | Paired Student's *t* test or Wilcoxon signed rank test | na | **GBZ effect on early:**<br>L2/3 sink p=0.013<br>L5 sink p=0.25<br>L2/3 source p=0.026<br>L5 source p=0.19 |
| Suppl. 6D | Paired Student's *t* test or Wilcoxon signed rank test | na | **GBZ effect on late:**<br>L2/3 sink p=0.006<br>L5 sink p=0.00012<br>L2/3 source p=0.00006<br>L5 source p=0.0001 |
| Suppl. 6E | Paired Student's *t* test or Wilcoxon signed rank test | na | **GBZ effect on delayed:**<br>L2/3 sink $p=6x10^{-5}$<br>L5 sink $p=6x10^{-7}$<br>L2/3 source $p=6x10^{-5}$<br>L5 source $p=6x10^{-5}$ |
| Suppl. 7C | Paired Student's *t* test or Wilcoxon signed rank test | na | **GBZ effect on early:**<br>L2/3 sink p=0.16<br>L5 sink p=0.38<br>L2/3 source p=0.14<br>L5 source p=0.08 |
| Suppl. 7D | Paired Student's *t* test or Wilcoxon signed rank test | na | **GBZ effect on late:**<br>L2/3 sink p=0.0023<br>L5 sink p=0.0011<br>L2/3 source p=0.008<br>L5 source p=0.0005 |
| Suppl. 7E | Paired Student's *t* test or Wilcoxon signed rank test | na | **GBZ effect on delayed:**<br>L2/3 sink p=0.002<br>L5 sink p=0.006<br>L2/3 source p=0.002<br>L5 source p=0.0004 |
| 4E | Paired Student's *t* tests | na | **Early vs Late CSD peak**<br>L2/3 source: p = 0.18<br>L2/3 sink: p = 0.11<br>L5 source: p = 0.08<br>L5 sink: p = 0.86 |
| Suppl. 8E | Friedman rank sum test | Wilcoxon signed rank tests, Šidák correction, α=0.017 | Friedman: $p=2.2x10^{-16}$<br>Baseline vs Initial: $p=4.3x10^{-13}$<br>Baseline vs Reduction: p=0.2<br>Baseline vs Delayed: $p=1.6x10^{-7}$ |
